# Induction of Sertoli cells from human fibroblasts by NR5A1 and GATA4

**DOI:** 10.1101/430900

**Authors:** Jianlin Liang, Nan Wang, Jing He, Jian Du, Yahui Guo, Lin Li, Kehkooi Kee

## Abstract

Sertoli cells are essential nurse cells in the testis that regulate the process of spermatogenesis and establish the immune-privileged environment of the blood-testis-barrier (BTB). The induction of human Sertoli cells from fibroblasts could provide cellular sources for fertility and transplantation treatments. Here, we report the *in vitro* reprogramming of human fibroblasts to Sertoli cells and characterize these human induced Sertoli cells (hiSCs). Initially, five transcriptional factors (NR5A1, GATA4, WT1, SOX9 and DMRT1) and a gene reporter carrying the AMH promoter were utilized to obtain the hiSCs. We further reduce the number of reprogramming factors to two, i.e., NR5A1 and GATA4, and show that these hiSCs have transcriptome profiles that are similar to those of primary human Sertoli cells. Consistent with the known cellular properties of Sertoli cells, hiSCs attract endothelial cells and exhibit high number of lipid droplets in the cytoplasm. More importantly, hiSCs can sustain the viability of spermatogonia cells harvested from mouse seminiferous tubules. In addition, hiSCs suppress the production of IL-2 and proliferation of human T lymphocytes. When hiSCs were cotransplanted with human embryonic kidney cells, these xenotransplanted human cells survived longer in mice with normal immune systems. hiSCs also allow us to determine a gene associated with Sertoli-only syndrome (SCO), *CX43*, is indeed important in regulating the maturation of Sertoli cells.

## Introduction

Sertoli cells are the first somatic cell type to differentiate in the testis and the only somatic cell type inside the seminiferous tubules. Sertoli cells play a critical role in directing testis morphogenesis and the creation of an immune-privileged microenvironment, which is required for male germ cell development. During early gonad development, male somatic cells express the male sex-determining gene *SRY*, which directs the sex-specific vascular development and seminiferous cord formation ^1–3^ via the initiation of a cascade of genes, including *SOX9, FGF9, AMH* and *PGD2* ^4,5^. *NR5A1* (or *SF1*)*, GATA4*, and *WT1* are major transcriptional factors that direct somatic cells to become fetal Sertoli cells ^6^. The expanding fetal Sertoli cells and another type of testicular somatic cell (i.e., peritubular cells) regulate the final organization and morphogenesis of the developing gonad into a testis ^7,8^.

Sertoli cells are the pivotal somatic cell regulators inside the seminiferous cord. Sertoli cells embed male germ cells during all differentiating stages and provide immunological, nutritional and structural support for germ cell development ^9^. Sertoli cells secrete the growth factors and cytokines needed for proper spermatogenesis, including the maintenance of spermatogonial stem cells, meiosis initiation of spermatocytes, and maturation of spermatozoa ^10^. Furthermore, Sertoli cells have the unique ability to modulate immunoreactions that protect the developing germ cells from immunological attacks. The immune-privileged potential of Sertoli cells has been utilized in many allo- and xeno-grafts to reduce the immune response in the field of cell transplantation ^11–13^. Preclinical studies have transplanted Sertoli cells with various other cell types for the treatment of diabetes, neurodegenerative diseases, Duchenne muscular dystrophy, skin allografts and other diseases ^14^.

Recently, co-cultures of differentiated rodent primordial germ cells and neonatal testicular somatic cells have successfully enabled meiosis completion and round spermatid formation *in vitro* ^15^, highlighting the potential use of testicular somatic cells in the field of reproductive medicine. Human pluripotent stem cells have been differentiated to spermatid-like cells ^16,17^, but the co-culturing of stem cells with Sertoli cells could enhance the efficiencies of obtaining functional male gametes. However, the procurement of human Sertoli cells is not feasible because of biological and ethical constraints. The availability of donated Sertoli cells is limited, and expanding the limited number of human Sertoli cells *in vitro* remains a challenge ^18,19^. Therefore, the generation of patient-specific Sertoli cells from fibroblasts could alleviate these issues and fulfill the basic research and clinical demands.

Direct lineage reprogramming has been considered a promising strategy for obtaining functional cell types with lower teratoma risks than directed differentiation of pluripotent stem cells ^20,21^. The induction of cell type conversion between divergent lineages has been achieved using combinations of lineage-specific transcription factors ^22–25^. Fibroblasts are common cells in animal connective tissues that can be conveniently obtained from patients. Therefore, fibroblasts are often used as initiating cells in many lineage reprogramming experiments. The direct reprogramming of Sertoli cells from fibroblasts has been demonstrated in mouse by overexpressing five defined transcriptional factors ^26^, but the direct lineage conversion of human Sertoli cells from fibroblasts has not been described. Here, we report the efficient induction of human Sertoli cells (hiSCs) from both human pulmonary fibroblasts (HPF) and fibroblasts derived from human embryonic stem cells (hESCs) by overexpressing GATA4 and NR5A1. These hiSCs exhibit an epithelial morphology, lipid droplet accumulation, and transcriptomes similar to those of primary Sertoli cells; sustain the growth of mouse spermatogonia cells; and perform immune-privileged function during transplantation experiments.

Connexin 43 (CX43) is a predominant gap junction protein expressed in BTBs that affects the maturation of Sertoli cells and spermatogenesis ^27–30^. The deletion of CX43 in Sertoli cells, but not germ cells, causes infertility in mice ^27,31^. The absence of CX43 expression in human Sertoli cells is associated with Sertoli cell-only syndrome (SCO) and impaired spermatogenesis in male patients ^32,33^, but whether the deletion of *CX43* directly affects the characteristics of human Sertoli cells has not been demonstrated. Utilizing our *in vitro* hiSC model, we demonstrate that the deletion of *CX43* affects the transcriptome profile and maturation of hiSCs.

## Results

### Testing the reprogramming capability of five putative transcriptional factors

Based on the reprogramming capability of the transcriptional factors reported in a mouse study ^26^, we first tested the reprogramming capabilities of the human homologs of the five transcriptional factors (5TFs: NR5A1, GATA4, WT1, SOX9 and DMRT1) to convert human fibroblasts to hiSCs. All five human homologs were correctly cloned into lentiviral vectors and expressed at high levels as verified by immunofluorescent staining of HPFs (Fig. 1a). After the lentiviral transduction with all five factors and culturing in selective medium for 5 days, many HPFs started to transform from the typical elongated morphology of fibroblasts into the squamous morphology that typically appears in epithelial cells (Supplementary Fig. 1a). The analysis of the transcriptional expression showed that genes enriched in Sertoli cells, such as *AR, KRT18, CLU, PTGDS, SCF, BMP4 and INHA*, exhibited increased expression in this mixed population of transformed HPFs (Supplementary Fig. 1b). To determine whether these 5TFs were able to reprogram other fibroblast sources, we derived human fibroblast-like cells from hESC line H1 (dH1) and reprogrammed these cells as described for the HPFs. The dH1 morphology resembled that of fibroblasts, and no detectable expression of pluripotent markers was observed, but the expression of many markers of fibroblasts was observed (Supplementary Fig. 2a,b,c). After the transduction of the 5TFs, dH1 underwent a fibroblast to epithelial transformation similar to that observed in the HPFs (Supplementary Fig. 2d), suggesting that the 5TFs can transform both types of fibroblasts into epithelial-like cells and increase the expression of Sertoli cell markers.

**Figure 1.**
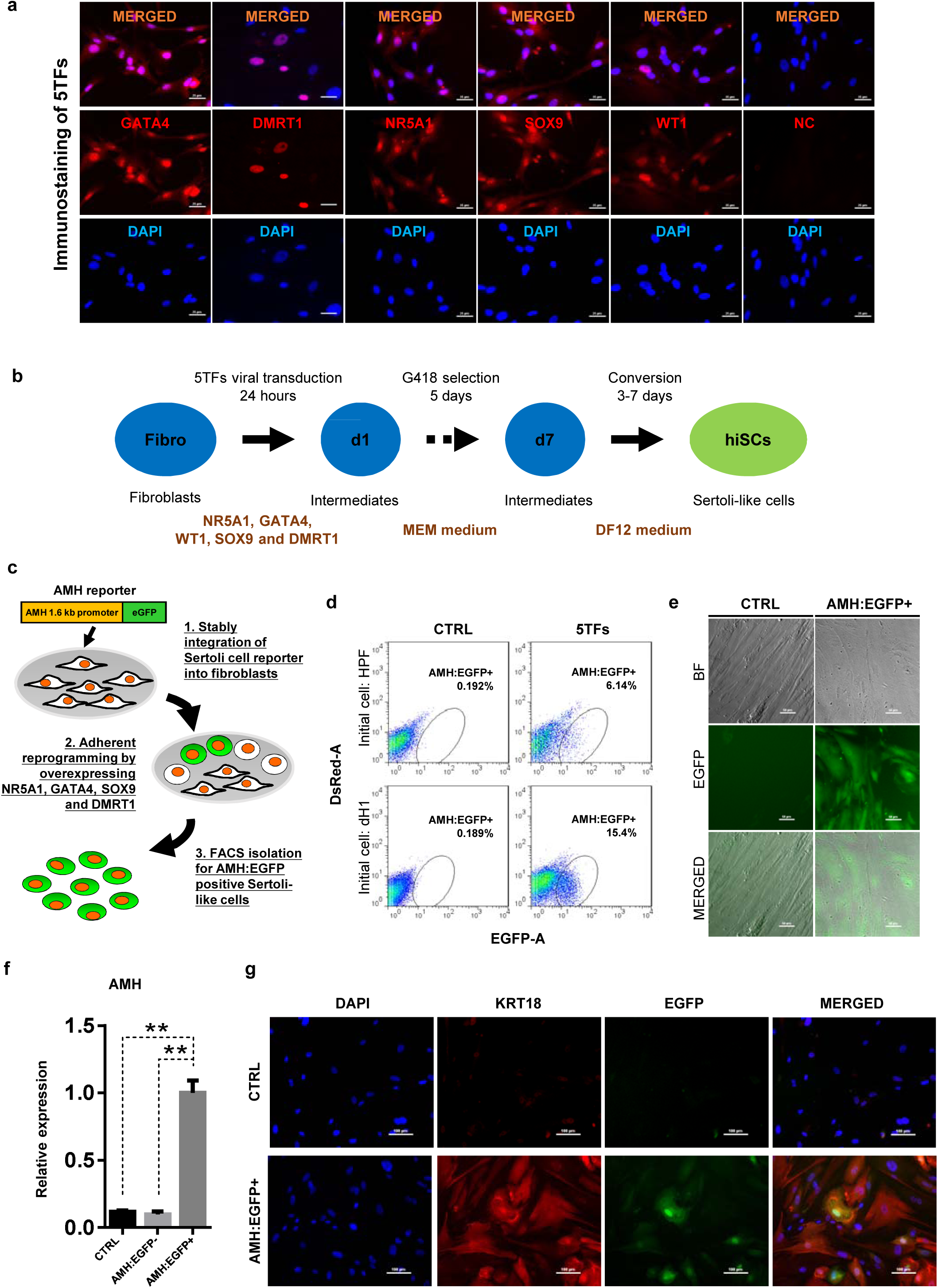
Induction of Sertoli-like cells (hiSCs) from human fibroblasts. (a) Immunostaining of NR5A1, GATA4, WT1, SOX9 and DMRT1 in human fibroblasts (HPF) 3 days post infection. DAPI was used to indicate nucleus. Scale bar, 25 μm. (b) Experimental design for the reprograming of human Sertloli-like cells (hiSCs). Human fibroblasts were infected with lentivirus carrying human transcription factors: NR5A1, GATA4, WT1, SOX9 and DMRT1 (5TFs, Table S1). The culture medium was changed to MEF medium 1 day after infection and followed by G418 selection for 5 days, then changed to DFMEM/F12 medium. hiSCs were characterized 10–14 days post viral transduction. (c) Schematic protocol for the reprogramming and isolation of AMH positive Sertoli-like cells. An AMH:EGFP reporter was integrated to the fibroblasts for cell sorting. (d) FACS analysis of AMH:EGFP+ cell at day 10 after 5TFs infection, the percentage of EGFP+ cell was used to determine the induction efficiency. Two types of human fibroblasts (HPF and dH1) were tested. Cells infected by p2k7 empty vector were used as negative control (CTRL). (e) Morphology of AMH:EGFP+ hiSCs after FACS. Scale bars, 20 μm. (f) The mRNA level of *AMH* was enriched in AMH:EGFP+ hiSCs compared to the control group as measured by quantitative PCR. n=3, technical replicates of ~5 × 10^4^ cells were collected by FACS. All data are presented as means ± SD. *p < 0.01; **p < 0.05. > 3 independent experiments were carried out. (g) Immunofluorescent staining of KRT18 in AMH:EGFP+ hiSCs and dH1 infected by p2k7 empty virus (CTRL) after FACS. Scale bar, 50 μm.

To isolate and enrich the hiSCs in the mixed population of reprogrammed fibroblasts, we constructed a gene reporter system utilizing the 1.6 kb promoter region of *AMH*, which is a gene specifically expressed in Sertoli cells ^34^, connected to EGFP (Fig. 1c). When the lentivector carrying the reporter, AMH:EGFP, was stably integrated into HPFs and dH1, none of the cells would express EGFP without transcriptional induction. After the transduction and selection of the fibroblasts, some reprogrammed cells were expected to express EGFP to allow us to isolate them by fluorescence-activated cell sorting (FACS). We found that both HPF and dH1 reproducibly yielded a clear AMH:EGFP-positive population after 10 days of transduction with the 5TFs (Fig. 1d). Intriguingly, the AMH:EGFP+ population in the dH1 group (~15%) was much higher than that in the HPF group (~6%), indicating that the conversion of the dH1 cells was more efficient. The AMH:EGFP+ cells were isolated by FACS and adhered to culture dishes and exhibited an epithelial morphology (Fig. 1e). We verified that the endogenous AMH expression was activated because the expression level of the *AMH* gene was significantly upregulated in the AMH:EGFP+ cell population compared to that in the control dH1 cells (CTRL) and AMH:EGFP- cells (Fig. 1f). Moreover, an epithelial marker expressed in Sertoli cells, i.e., KRT18, was used to validate the transformation of the fibroblasts to Sertoli cells. The cytoskeleton pattern of KRT18 expression was observed in the AMH:EGFP+ cells but not the control dH1 cells (Fig. 1g).

### Whole-genome transcriptional profiling of hiSCs resembling adult Sertoli cells

To determine whether hiSCs reprogrammed with the 5TFs (5F-hiSCs) are similar to human Sertoli cells, we compared the transcriptomes of the AMH:EGFP+ 5F-hiSCs, dH1 cells infected with p2k7 empty virus (dH1-2K7) as negative controls, and primary adult Sertoli cells (aSCs) from human biopsy samples. We focused our analysis on the differentially expressed genes (DEGs, > 2-fold change, *p-value* < 0.05) between 5F-hiSCs and dH1-2K7 and between aSCs and dH1-2K7. In total, 7533 genes were differentially expressed between 5F-hiSCs and dH1-2K7, including 4528 upregulated genes and 3005 down-regulated genes (Fig. 2a,b). Additionally, 5377 genes were differentially expressed between aSCs and dH1-2K7, including 3343 upregulated genes and 2034 down-regulated genes (Fig. 2a, b). The Venn analysis showed that 3626 genes were shared among the DEGs in both hiSCs and aSCs, accounting for approximately 67% of the DEGs in aSCs. Among this shared group of DEGs (CO-DEGs), 1973 genes were upregulated, while 1314 genes were down-regulated in both the hiSCs and aSCs (Fig. 2c), indicating that the trends in transcriptional expression between the hiSCs and aSCs were the same in these genes. The cluster analysis of dH1-2K7, hiSCs and aSCs also showed that the CO-DEGs had a similar expression pattern between the hiSCs and aSCs, and consistency was observed between duplicate samples (Fig. 2d). The Gene Ontology (GO) analysis of the CO-DEGs showed that among the 1973 upregulated genes, many genes were involved in the regulation of cell communication, regulation of immune response processes, response to hormones, and lipid metabolic process, whereas among the 1314 down-regulated genes, many genes were involved in the mitotic cell cycle and microtubule-based processes (Fig. 2e). These changes in gene expression indicated that the hiSCs acquired unique cellular characteristics that were distinct from the original fibroblasts. To further confirm that the AMH:EGFP+ 5F-hiSCs have the signature of Sertoli cells, we examined the expression of several Sertoli cell markers, including *CLU, NCAM2, DHH, ERBB4, INHB, INHA, SHBG, GATA6, CDKN1B, TGFα* and *LMMA3*, by quantitative PCR (qPCR). Compared to the control cells, all Sertoli cell markers were highly enriched in the AMH:EGFP+ 5F-hiSCs (Fig. 2f). Taken together, the transcriptional profile of the AMH:EGFP+ 5F-hiSCs resembled that of the aSCs, and many Sertoli cell markers were expressed.

**Figure 2.**
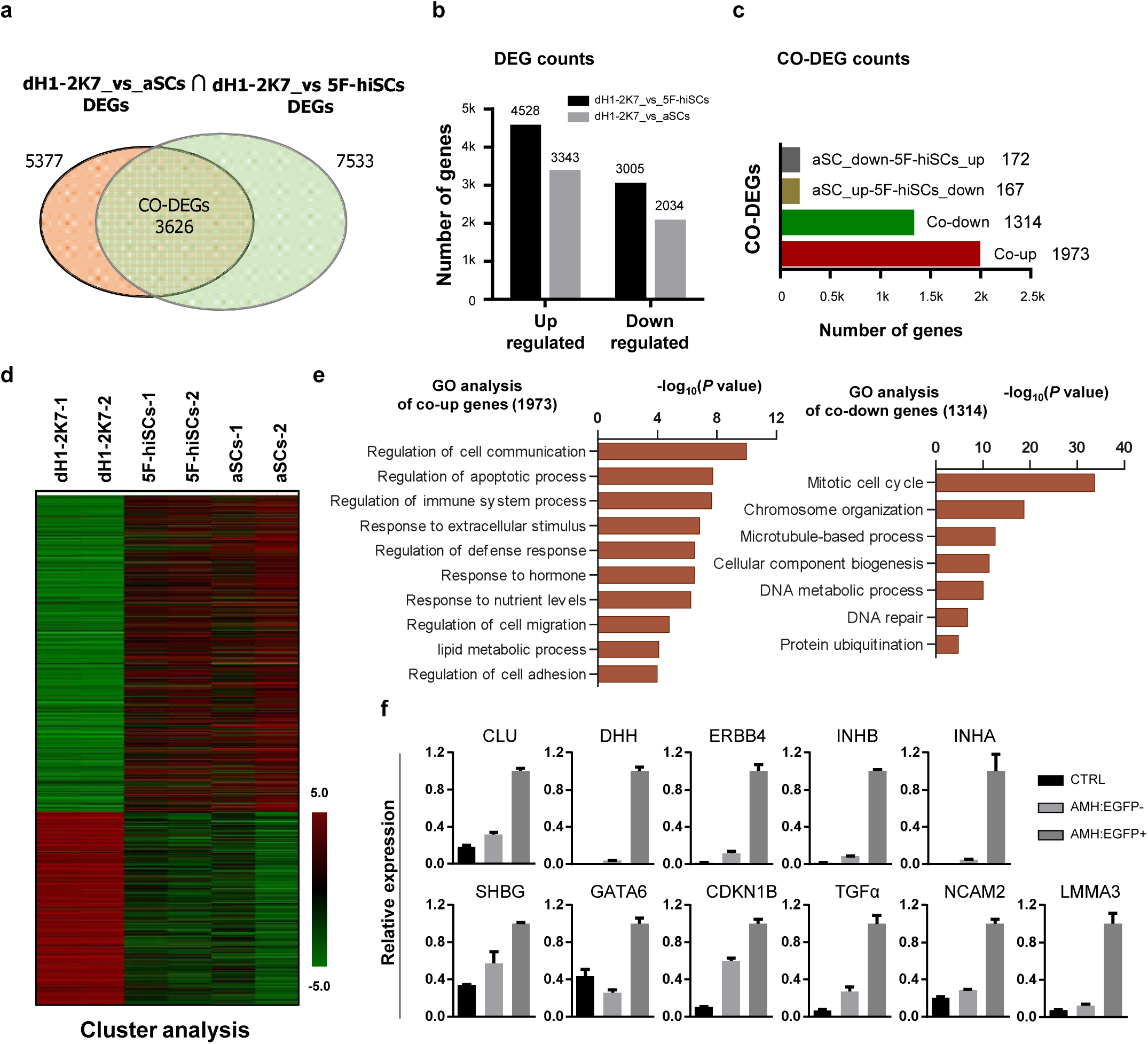
Whole genome transcriptional profiling of AMH:EGFP+ hiSCs. (a) Venn diagram to display differentially expressed genes (DEGs, FPKM value, fold change > 2) between dH1-2K7_vs_aSCs (n=2) and dH1-2K7_vs_5F-hiSCs (n=2). Intersection part represented the number of co-differentially expressed genes (CO-DEGs). (b) Upregulated and downregulated DEGs in (a). X axis indicates upregulated DEGs and downregulated DEGs. Y axis represented DEGs numbers. Comparing samples were shown by different colors. Black bar represented dH1-2K7_vs_5F-hiSCs. Gray bar represented dH1-2K7_vs_aSCs. (c) Upregulated and downregulated CO-DEGs in (a). X axis indicates CO-DEGs numbers. Red bar indicates co-upregulated DEGs, green bar indicates co-downregulated DEGs. Brown and gray bar indicate CO-DEGs in opposite trend of upregulation and downregulation respectively. (d) Heat map of gene expression of dH1-2K7, 5F-hiSCs and aSCs (n=2, two independent samples were subjected to RNA-seq). Red indicated upregulated expression, green indicated down-regulated expression. (e) Functional enrichment analysis, biological processes of 1973 and 1314 differentially expressed genes were showed. (f) The mRNA level of Sertoli-cell markers in AMH:EGFP-cells and AMH:EGFP+ cells as measured by qPCR. Fold expression was normalized to the levels of dH1 infected by empty virus p2k7 (CTRL). GAPDH was used as the housekeeping gene for normalization. All data are presented as means ± SD, n=2, 3 independent experiments were carried out.

### NR5A1 and GATA4 are sufficient to reprogram fibroblasts to hiSCs as 5F-hiSCs

Although the 5TFs yielded the AMH:EGFP+ hiSCs, the combination of all 5TFs may not be necessary to reprogram fibroblasts to hiSCs. Therefore, we used fewer transcription factors to generate hiSCs and compared the percentage of AMH:EGFP+ cells in all 31 combinations of NR5A1, GATA4, SOX9, WT1 and DMRT1 (Table S2). The FACS results indicated that 16 combinations of transcriptional factors yielded varying levels of AMH:EGFP+ cells after 10 days of reprogramming (Fig. 3a). NR5A1 was the only common factor found in all 16 combinations in AMH:EGFP+ cells. This transcriptional factor alone generated approximately 3.79% of AMH:EGFP+ cells, and the combination of all 5TFs produced the highest percentage of AMH:EGFP+ cells. Surprisingly, the combinations with NR5A1 and GATA4 generated as many AMH:EGFP+ cells as all 5TF combined. Moreover, all combinations containing NR5A1 and GATA4 resulted in a similar level higher than the combinations with NR5A1 (Fig. 3b). The AMH:EGFP+ cells generated by 2TFs and 5TFs showed similar morphologies, including a large cell body with an epithelial morphology, and expressed KRT18 (Fig. 1g,3c).

**Figure 3.**
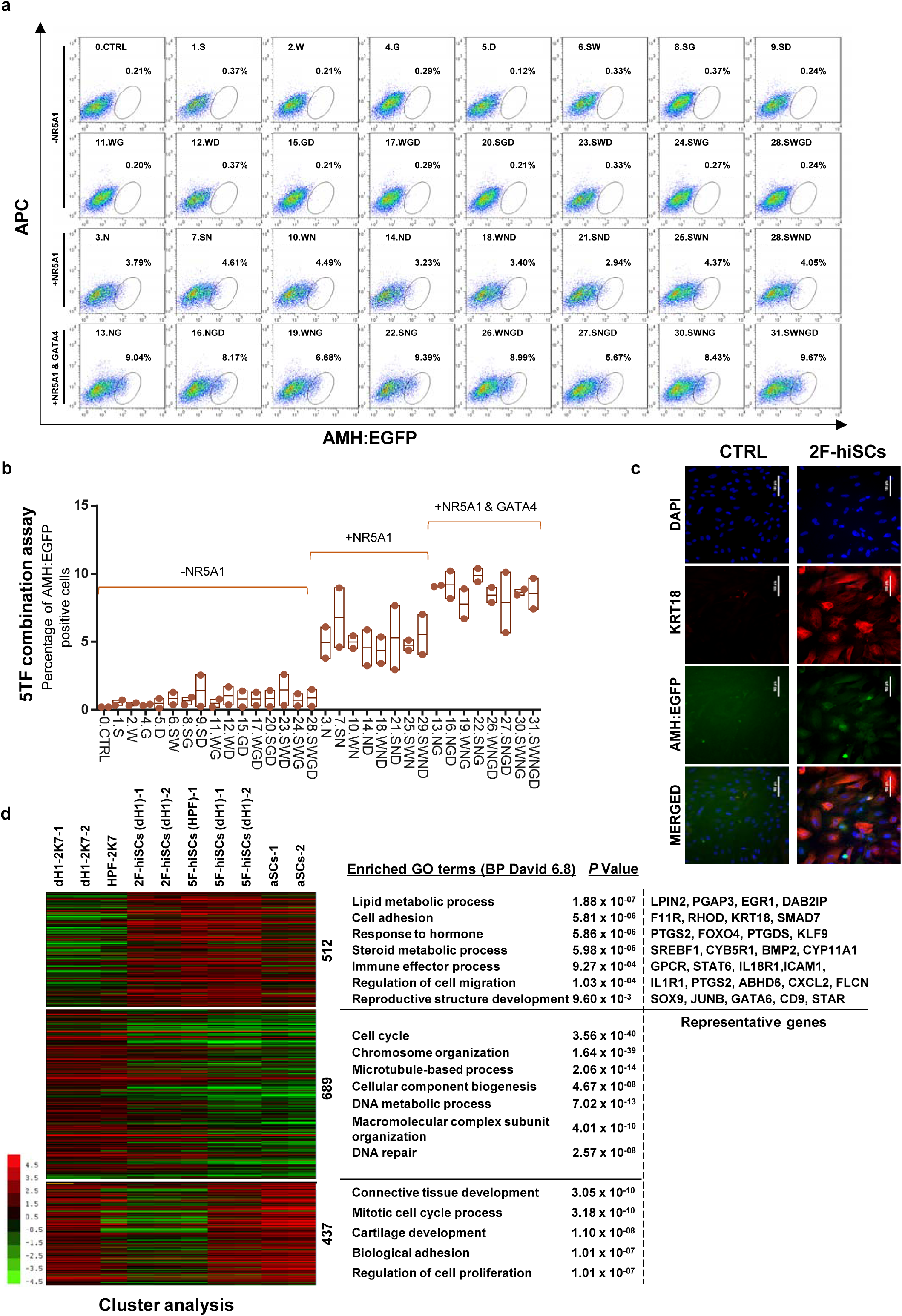
NR5A1 and GATA4 are sufficient to derive hiSCs. (a) Representative FACS results of different combinations of NR5A1, GATA4, WT1, SOX9 and DMRT1 for the induction of AMH:EGFP+ cells. dH1 fibroblasts were transduced with the indicated factors and reprogrammed for 10 days. The combinations were divided into three groups: −NR5A1, combinations without NR5A1; +NR5A1, combinations with NR5A1 but without GATA4; +NR5A1 & GATA4, combinations with both NR5A1 and GATA4. (b) Quantitative data of EGFP+ cells in (a) (n=2, biological replicates, error bar indicates SD), 3 independent combination experiments were conducted. ~10^4^ cells were FACS in each experiment. (c) Immunofluorescent staining of KRT18 in 2F-hiSCs and dH1-2K7 (CTRL) after FACS. Scale bar, 100 μm. (d) Heat map of RNA-seq data illustrating differentially co-expressed genes. Red indicates upregulated expression, whereas green indicates downregulated expression. The genes were divided into three groups: Genes upregulated in 2F-hiSCs (dH1), 5F-hiSCs (dH1) and aSCs; genes downregulated in 2F-hiSCs (dH1), 5F-hiSCs (dH1) and aSCs; and genes varying among different groups. The gene number of each group was listed next to the map. Functional enrichment terms of each group and the representative genes were shown on the right side of heat map for the upregulated gene group.

Then, we analyzed and compared the transcriptome of the 2F-hiSCs and with the transcriptome of the aSCs and 5F-hiSCs. We identified the common DEGs among the dH1/aSCs, dH1/5F-hiSCs (dH1), dH1/2F-hiSCs (dH1) and dH1/5F-hiSCs (HPF) and performed hierarchical clustering using their FPKM values. The analysis revealed that the transcriptome profiles of the 2F-hiSCs were more similar to the profiles of the 5F-hiSCs and adult Sertoli cells (aSCs) than to the dH1 and HPF profiles (data not shown). To identify the putative signature genes similar among the 2F-hiSCs, 5F-hiSCs and aSCs, we generated a heat map of the 1638 CO-DEGs and carried out gene correlation clustering. Notably, the differentially expressed genes were grouped into three groups of 512, 689 and 437 genes. The Gene Ontology analysis showed that many of the 512 highly expressed genes mostly shared by the 2F-hiSCs, 5F-hiSCs and aSCs were involved in reproductive structure development, immune effector processes and response to hormones. These genes, including *KRT18, PTGDS* and *SOX9*, are used as markers of Sertoli cells or are highly expressed in Sertoli cells ^26,35^. We further compared the expression of 59 genes that are highly expressed in Sertoli cells ^36–38^ among the 2F-hiSCs, 5F-hiSCs and aSCs. All three groups exhibited similar expression of many markers that are expressed in more mature Sertoli cells, including *CDKN1B*(or *p27^kip1^*) and *CLU*^39^. However, some markers in the aSCs, including *NCAM2, INHA* and *KRT18* ^39,40^, were expressed at lower levels (Supplementary Fig. 3) than those in the other two hiSCs. The Sertoli cell marker expression was very similar between the 2F-hiSCs and 5F-hiSCs. Therefore, we focused on the 2F-hiSCs for the subsequent more thorough characterization.

### 2F-hiSCs attract human endothelial cells and accumulate lipid droplets

Sertoli cells mediate the migration of endothelial cells to seminiferous tubules during testicular cord formation ^3,41^. We investigated whether the hiSCs attracted human umbilical vein endothelial cells (HUVECs). The migration assay showed that compared to ~40 cells under control conditions, the number of HUVECs that were attracted and passed through the membrane was ~80 cells per unit after 20 hours of induction in conditioned media. These results indicate that significantly more endothelial cells were attracted by the conditioned medium collected from the 2F-hiSCs than by that collected from the dH1-2K7 cells (Fig. 4a,b).

**Figure 4.**
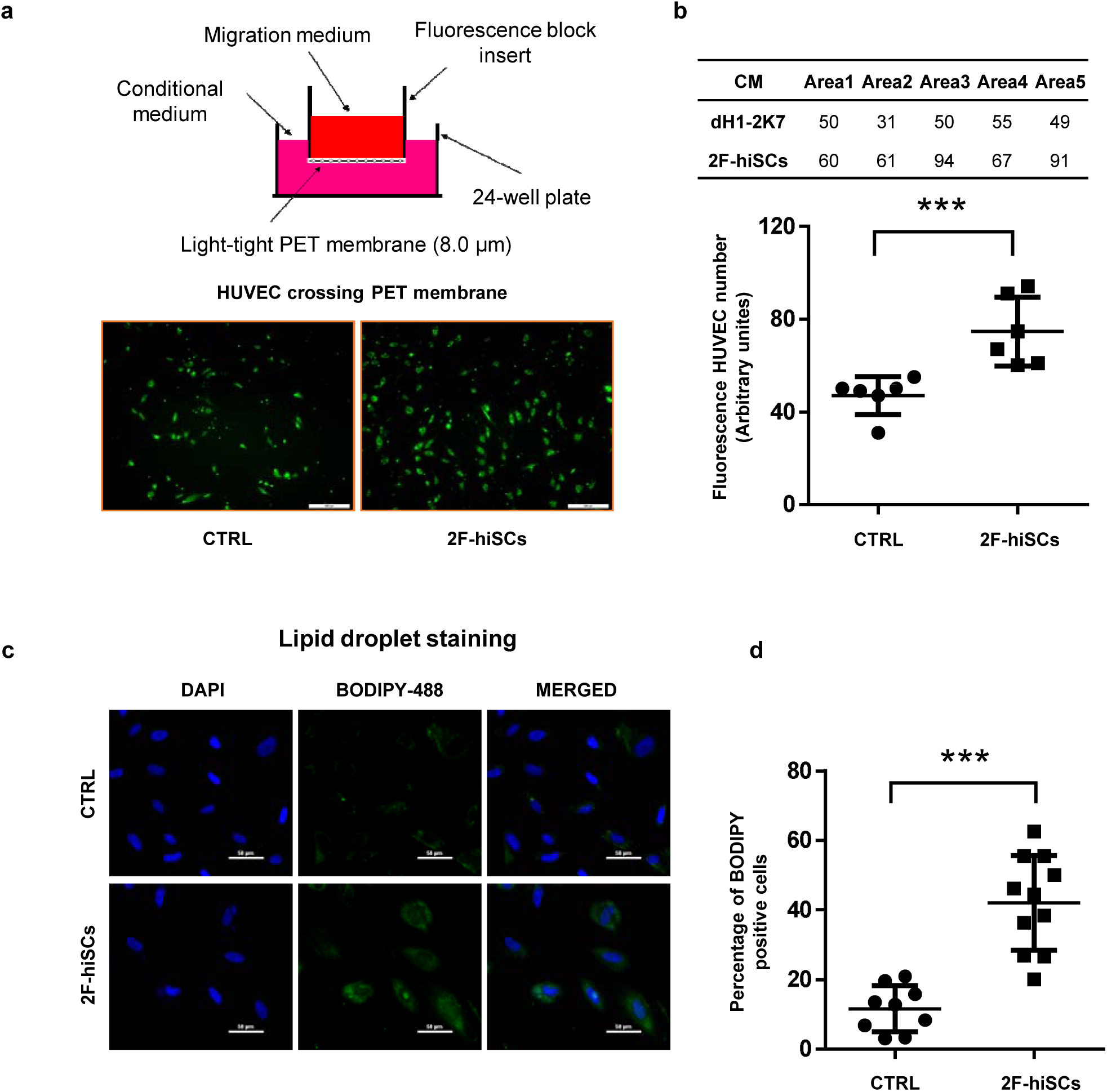
2F-hiSCs attract endothelial cells and accumulate lipid droplets. (a) Migration assay of HUVECs incubated with the indicated conditioned media for 20 hours. HUVEC were treated with 2.5 μM Calcein-AM fluorescent dye 1 hour prior to seeding. Top, experimental diagram using the fluoroblok 24-multiwell insert plates with 8.0 μm pores. Down, fluorescent images showing cells passed through the pores. (b) Number of HUVEC cells crossing through the transwell membrane. Each data point on the graph indicates the number of cells in one independent area. Scar bar, 300 μm. Error bars indicate standard deviations of all replicates. (c) BODIPY staining of lipid droplets in dH1-2K7 (CTRL) and 2F-hiSCs. All cells were fixed with 4% PFA and then stained with BODIPY for lipid droplets and DAPI for nucleus. (d) Each data point on the graph indicates the number of strong BODIPY+ cells in one image area. Scar bar, 50 μm. Error bars indicate standard deviations of all replicates.

Another unique feature of Sertoli cells in humans and other mammalian species is the accumulation of lipid droplets in the cytoplasm ^42,43^. The Gene Ontology analysis showed that genes that participate in lipid metabolism processes were upregulated in the 2F-hiSCs, 5F-hiSCs and aSCs (Fig. 3d), supporting the presence of high numbers of lipid droplets in the hiSCs. We used BODIPY to stain and count the number of cells with high numbers of lipid droplets to determine whether lipid droplets appeared in the 2F-hiSCs. The average percentage of cells that exhibited strong BODIPY positivity was approximately 15% in the control cells and approximately 40% in the 2F-hiSCs. Therefore, the percentage of cells containing high quantities of lipid droplets was approximately 2.7-fold higher in the 2F-hiSCs group than that in the dH1-2K7 group (Fig. 4c, d)

### 2F-hiSCs sustain *in vitro* culturing of mouse spermatogonia cells

Mouse spermatogonia cells were isolated from seminiferous tubules of 6 dpp mice and co-cultured with dH1 or 2F-hiSCs to examine whether 2F-hiSCs sustain the growth of male germ cells (Fig. 5a, Supplementary Fig. 4). More mouse germ cells attached and survived on the 2F-hiSCs than dH1 cells after a 12-hour culture (Fig. 5b). The morphology of the round cells attached to the hiSCs appeared alive and resembled spermatogonia cells. The cells attached to dH1 appeared apoptotic and degenerated. The co-cultured samples were fixed and immunostained 48 hours after plating to further confirm that the 2F-hiSCs formed solid attachments with the spermatogonia cells. The immunostaining of the germ cell-specific marker DAZL (to identify mouse spermatogonia cells) and human-nuclear specific marker NuMA (to identify hiSCs and dH1) indicated that significantly more DAZL-positive spermatogonia cells attached to the hiSCs, but almost no DAZL-positive cells attached to the dH1 cells despite the similar numbers of plated hiSCs and dH1 cells (Fig. 5c,d). Sertoli cells directly contact male germ cells in seminiferous tubules *in vivo*, and we hypothesized that the hiSCs would directly contact the spermatogonia cells. We immunostained the co-cultured cells with DAZL and KRT18 and found that many DAZL-positive cells localized to areas occupied by hiSCs, typically at the edge of the cell bodies (Fig. 5e).

**Figure 5.**
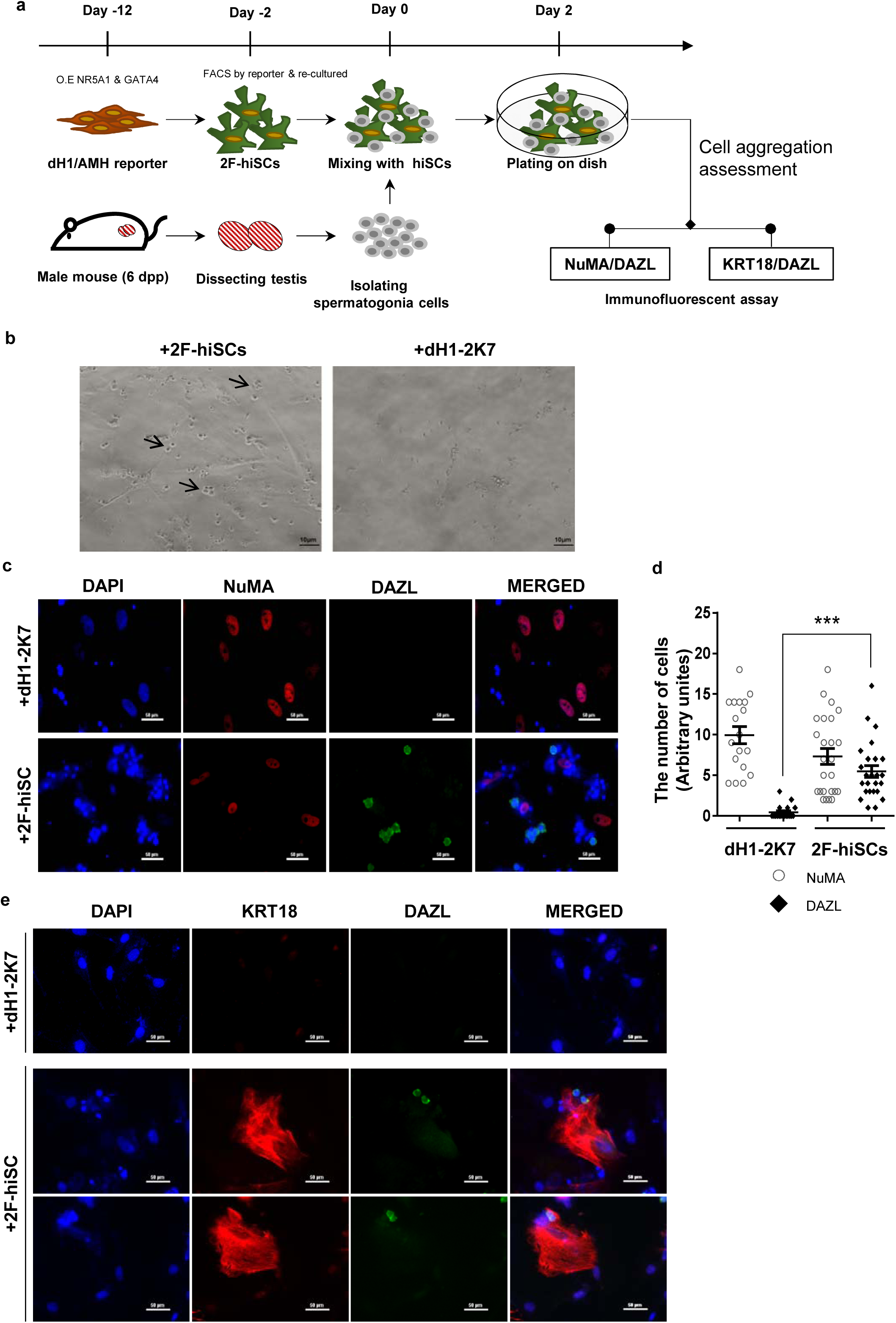
2F-hiSCs sustain the viability of mouse spermatogonia cells. (a) Timeline and steps of co-culturing hiSCs with mouse germ cells. (b) Mouse spermatogonia cells plated on 2F-hiSCs or dH1-2K7. Bright-field picture was taken after 12 hours of plating. More round and healthy cells (indicated with the arrow heads) attached to hiSCs than dH1-2K7 cells. Scale bar, 10 μm. (c) Immunofluorescent staining of germ cell marker, DAZL (green), and human-cell-specific marker, NuMA (red), in co-cultured mouse spermatogonia cells with either 2F-hiSCs or dH1-2K7, 48 hours after plating. DAPI (blue) was used to indicate the nuclei. Scale bar, 50 μm. (d) Cell numbers with indicated markers on the co-cultured plate in (c). Each data point indicates the number of cell counted in one image area. Error bars are the standard deviations of all the counted data points. (e) Immunofluorescent staining of DAZL (green) and KRT18 (red) in co-cultured mouse spermatogonia cells with 2F-hiSCs, 48 hours after plating. DAPI (blue) was used to indicate the nuclei. Scale bar, 50 μm.

### 2F-hiSCs suppress T cell proliferation, IL-2 production and protect transplanted human cells

A specialized property of Sertoli cells is their functional role in creating an immune-privileged environment in seminiferous tubules to protect germ cells from immunological attacks. Previous studies have shown this unique function, which has been exploited in therapeutic transplantation for the protection of many other cell types ^11^. We first investigated whether the medium from the 2F-hiSCs cells exhibited any suppressive effect on the proliferation of Jurkat E6 cells (human T lymphocytes) to examine whether the 2F-hiSCs could suppress the immunoreaction of immunological cells. The suppressive effect was evaluated using an assay commonly used to determine the active metabolism of a proliferating cell, i.e., WST-1^44^ (Fig. 6a). The Jurkat E6 cells were treated with various concentrations of 2F-hiSCs-conditioned media and exhibited a significant dose-responsive decrease in cell proliferation compared to that in cells treated with dH1-2K7-conditioned media (Fig. 6b). The proliferation level was ~35% lower in the Jurkat lymphocytes exposed to the highest concentration of the 2F-hiSC-conditioned medium. We collected the Jurkat cells and analyzed the protein levels of interleukin-2 (IL-2), which plays an essential role in the immune system. The ELISA indicated that the IL-2 levels were significantly lower in the cells cultured with the 2F-hiSCs-conditioned media than those in the cells cultured with the dH1-2K7-conditioned media (Fig. 6c). 21 gene previously known to participate in immunosuppression of Sertoli cells^12,45^ or categorized as immune effector gene by gene ontology were activated after reprogramming of dH1(Fig. 6d), further confirmed that hiSCs acquired immunosuppressive function. Approximately 1.3 × 10^6^ human 293FT cells stably integrated with a luciferase-expressing vector were co-transplanted with 2.5 × 10^5^ dH1-2K7 or 2F-hiSCs cells into mice with normal immune systems via hypodermic injection to determine whether the 2F-hiSCs could protect xenotransplanted cells. The transplantation experiment was performed to investigate the immunosuppressive effects at different locations (foreleg, hindleg, left and right sides of the animals as indicated on the figures) in two different animals, and the transplanted sites were monitored for up to 10 days. D-luciferin was injected into the animals to follow the surviving transplanted 293FT cells, and the signal was monitored using live imaging 15 minutes post-injection beginning 3 days after transplantation. The transplanted 293FT cells gradually diminished in the immunocompetent mouse from day 3 to day 10 (mouse #1, #2) as indicated by the reduced luciferase activity at the transplanted sites (Fig. 6e, Supplementary Fig. 5). All 293FT cells co-transplanted with hiSCs exhibited higher luciferase activity, which ranged from 1.7- to 3.9-fold, 3 days after transplantation. Three of the four groups of transplanted cells survived until day 10, and two of the three groups of 293FT cells with hiSCs survived at least 10 days after transplantation with strong luciferase activity. In contrast, their counterpart control cells exhibited less than 40-fold or no detectable luciferase activity (mouse #1 foreleg group and mouse #2 hindleg group).

**Figure 6.**
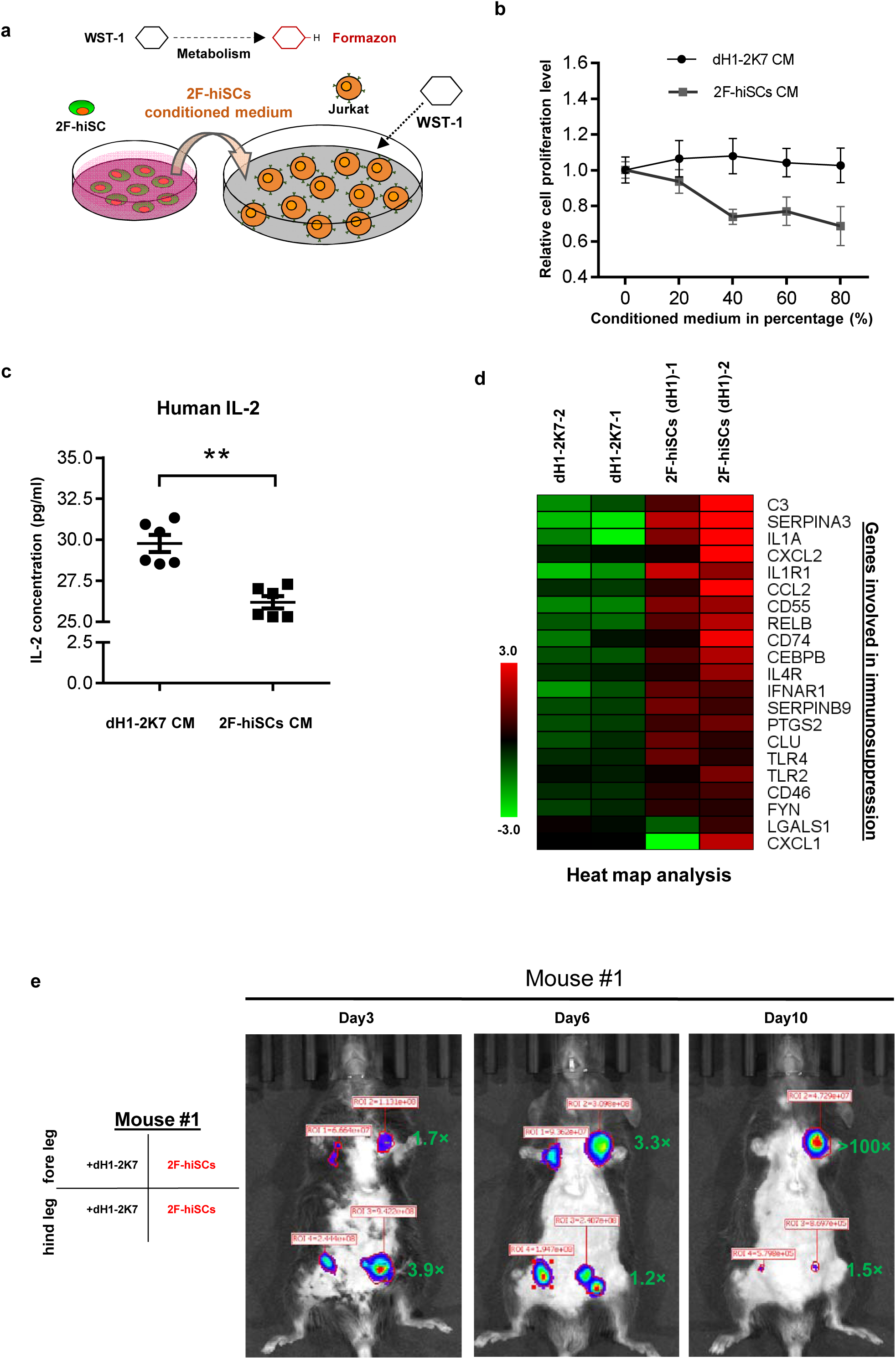
2F-hiSCs exhibit immunosuppressive function and protect human cells in xenotransplantations. (a) Schematic illustration of the WST-1 assay to measure the inhibiting effect of 2F-hiSCs on Jurkat-E6 cell proliferation. Cleavage of the tetrazolium salt WST-1 by metabolically active cells resulted in the formation of formazon. The level of formazon was directly proportional to the proliferation of the Jurkat cells. (b) Measurement of proliferation level in Jurkat cells treated with dH1-2K7 or 2F-hiSCs conditioned medium. X axis represent the indicated concentrations of conditioned medium. Conditioned medium from dH1-2K7 was used as a control. Values are average of triplicates, error bars indicates standard deviations, two separate experiments were conducted. (c) Histogram showing the production of human IL-2 in lymphocytes treated by dH1-2K7 or 2F-hiSCs conditioned medium, as measured by ELISA. Error bars indicates standard deviation of replicates from six samples. (d) Heat map analysis showing expression of genes involved in immunosuprression in control versus 2F-hiSCs. Genes are selected based on previous studies showing their involvement in immunosuppression of Sertoli cells. (e) Live imaging of luciferase-tracking assay 3 days to 10 days after transplantations. ~1.3 × 10^6^ 293FT cells were subcutaneously injected with either 2 × 10^5^ dH1-2K7 or 2F-hiSCs cells into the indicated sites of the mouse. Cell types and locations of transplantation are indicated at left. Number in red indicates primary readings of luciferase activity, green numbers indicate fold change of luciferase activity in hiSCs + 293FT cells normalized to the controls from day3 to day10 in mouse #1.

### CX43 deletion disrupts gap junctions and alters the expression profile of hiSCs

We investigated whether the deletion of the gap junction protein CX43 could affect hiSCs formation and determined whether hiSCs exhibit the same genetic requirements for development as Sertoli cells *in vivo*. We created a homozygous deletion of *CX43* hESC line, derived fibroblasts from this line and compared the reprogramming efficiency of the hiSCs (Fig. 7a). We successfully generated three targeted mutations at *CX43*, i.e., one single allele mutation (#6) and two double allele mutations (#21 & #26) at Exon 2 of the *CX43* gene, using a CRISPR-CAS9 gene editing system (Fig. 7b). The protein expression was completely disrupted in the CX43^-/-^ (#21) and CX43^-/-^ (#26) cell lines, but the heterozygous protein level remained similar to the wild-type CX43 level based on the Western blot analyses (Supplementary Fig. 6a). The immunostaining of CX43 (Fig. 7c) and photo bleaching assay (Fig. 7d,e) of ES-derived fibroblasts (dH1) both showed that the expression of CX43 and gap junctions between neighboring cells were disrupted because there was no detectable CX43 staining or diffusing fluorescent dye recovered after photo bleaching in the CX43^-/-^ (#26) cell line.

**Figure 7.**
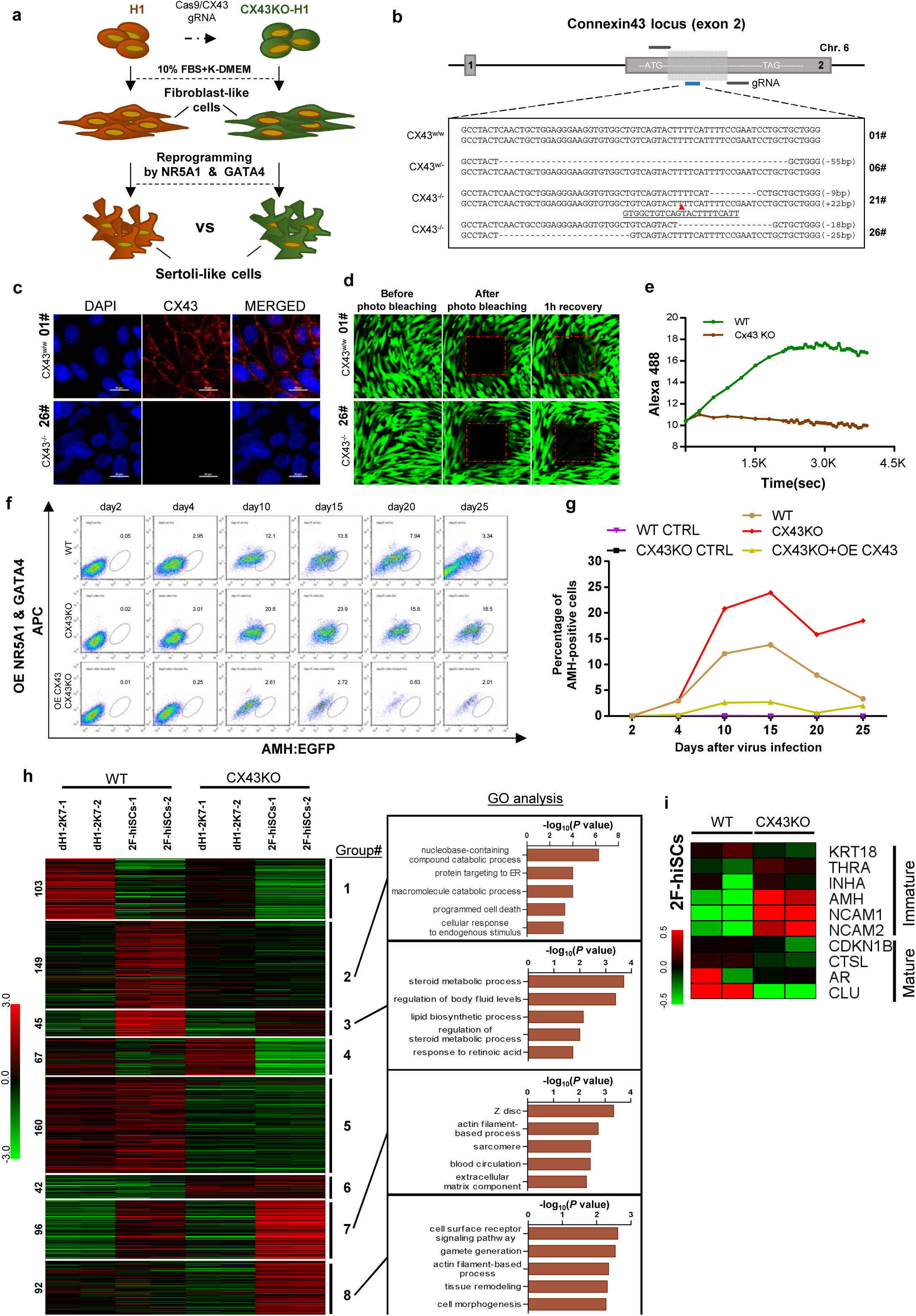
CX43 regulates maturation of Sertoli cells. (a) Schematic diagram showing the comparison of 2F-hiSCs cells reprogrammed from *CX43* knock out fibroblasts or wild type fibroblasts. CX43KO was initially carried out in H1 hESC lines. After confirmation of CX43 KO, the hESCs were differentiated to fibroblasts (dH1 or CX43KO dH1). The fibroblasts were then reprogrammed with NR5A1 and GATA4 into 2F-hiSCs. (b) *CX43* knockout design and the targeted sequences of the indicated hESC lines. The knockout region was located on the second exon of the *CX43* gene. Three knockout cell lines were correctly targeted and mutated, one was heterozygous (06#) and the other two were homozygous (21# and 26#). Dotted lines indicate deletion mutations, red triangles indicate insertion mutations. (c) Immunofluorescence analysis of CX43 (red) in wild type ES cell line (01#) and knock out cell lines (26#), DAPI (blue) was used to indicate the nuclei. Scale bar, 20 μm. (d) Photo bleaching assay to test the Calcien AM transport ability of fibroblasts generated from wild type ES cell line (01#) and knock out cell lines (26#). (e) Measurement of Calcien AM signal over time after fluorescence bleaching. (f) Time course experiment during 2F-hiSCs reprogramming. The AMH:EGFP+ percentage resulted from three initial fibroblasts were tested: wild type dH1, CX43 KO dH1 and CX43 KO plus CX43 overexpressed dH1. (g) Percentage of hiSCs(AMH:EGFP+) in WT, CX43KO and CX43KO+ CX43 overexpression from day 2 to day 25 of reprogramming. (h) Hierarchical clustering analysis using DEGs (FPKM value, fold change > 2) in Supplementary Fig. 7b. Genes were classified into 8 parts according to the expression pattern in CX43 KO dH1, CX43KO 2F-hiSCs, wild type dH1 and wild type 2F-hiSCs. Gene Ontology analysis of genes in part 2, 3, 7, 8 was shown on the right respectively. Other groups were shown in Supplementary Fig. 7c. (i) Heat map indicating expression level of immature and mature Sertoli-cell markers in WT and CX43KO 2F-hiSCs. Relative gene expression level was indicated as red (upregulated) or green (down-regulated).

We compared the reprogramming efficiency of the 2F-hiSCs between the wild-type dH1 and CX43KO (CX43^-/-^ (#26)) cell lines. The time course experiments showed that the percentage of AMH:EGFP+ in the WT hiSCs peaked at ~13.8% on day 15 and decreased to 3.3% on day 25 (Fig. 7f, g; Supplementary Fig. 6b). Remarkably, the percentage of AMH:EGFP+ in the CX43^-/-^ (#26) cell line was 23.9% on day 15 and decreased to 18.5% on day 25, but the overexpression of CX43 in the deletion cell line revealed a much lower percentage of AMH:EGFP+ from days 4 to 25. Therefore, the expression level of CX43 in the cells was indirectly proportional to the percentage of AMH:EGFP+ cells. This result is consistent with the higher AMH expression in CX43 knockout mice reported in a previous study ^30^ and suggests that the effect of the CX43 deletion leads to the dedifferentiation of Sertoli cells to a less mature state. The high percentage of AMH:EGFP+ cells in the CX43KO cells may be due to their more immature status than that of WT AMH:EGFP+ hiSCs. Therefore, we compared the transcriptional profiles of these two populations to examine whether CX43KO affected gene expression or any cellular processes. The volcano analysis revealed that 2736 genes were differentially expressed (p-value less than 0.01) between the CX43KO 2F-hiSCs and WT 2F-hiSCs (Supplementary Fig. 7a). We found 754 genes with a difference in the transcript level greater than two-fold; 512 genes were down-regulated and 242 genes were upregulated in the CX43KO 2F-hiSCs compared to those in the WT 2F-hiSCs (Supplementary Fig. 7b). We further analyzed the 755 genes using a heat map and GO analysis to identify specific genes or processes affected by CX43KO. The genes were classified into 8 gene sets according to the expression patterns among the WT dH1, WT 2F-hiSCs, CX43KO dH1 and CX43KO 2F-hiSCs (Fig. 7h, Supplementary Fig. 7c). The genes in Groups 1 and 4 exhibited lower expression levels in the WT 2F-hiSCs and CX43 2F-hiSCs than in the other two groups of control cells, suggesting that these genes were affected by the reprogramming process. The genes in Groups 5 and 6 exhibited similar patterns between the WT cells and CX43KO cells, suggesting that these two groups were affected by CX43KO. In contrast, the expression profiles of the genes in Groups 2, 3, 7, and 8 in the WT and CX43KO hiSCs differed from those in their counterpart controls and between the WT and CX43KO hiSCs. These genes may reflect the cellular maturation status of these two reprogrammed cell populations. The GO analysis of enriched terms in these 4 groups revealed that Group 2 was enriched with genes involved in catabolic processes, including nucleobase-containing compound catabolic process, and Group 3 contained genes involved in steroid metabolic or lipid biosynthetic processes. Group 7 included many genes that participated in the cytoskeleton, and Group 8 contained genes related to gamete generation.

Ten marker genes previously reported to be more highly expressed in mature or immature Sertoli cells ^35,39^ were chosen for the expression level comparisons between WT and CX43KO hiSCs. The CX43KO hiSCs exhibited a higher expression of some immature markers and lower expression of mature markers, suggesting that the CX43KO cells were more immature than the WT hiSCs.

## Discussion

This study reported that human fibroblasts can be reprogrammed to Sertoli-like cells using 5TFs (NR5A1, GATA4, WT1, DMRT1 and SOX9) or 2TFs (NR5A1 and GATA4). Fibroblasts from two sources, i.e., human pulmonary fibroblasts and fibroblasts derived from human embryonic stem cells, were reprogrammed to Sertoli-like cells and exhibited transcriptomes similar to those of primary adult Sertoli cells. The present study generated a Sertoli cell-specific gene reporter, i.e., AMH:EGFP, to enhance the efficiency of isolating a relatively pure population of hiSCs from a mixed population of cells at different reprogramming stages (Fig. 1, 2). AMH has been used in many conditioned knockout studies to specify the expression of Cre recombinase in Sertoli cells ^46^. We cloned the human promoter of *AMH* and fused it to EGFP and showed that the cell population expressing EGFP appeared only after reprogramming, and these cells expressed many Sertoli cell marker genes. The success of reprogramming human fibroblasts with the same factors used in a previous mouse study ^26^ confirms that the reprogramming capability of these 5TFs is conserved between humans and mice.

Another achievement of this study was the reduction in the reprogramming factors from 5TFs to 2TFs. NR5A1, which is often called SF1, was the most essential of the five factors because almost no AMH:EGFP+ cells appeared in the reprogramming experiments without NR5A1, even if GATA4, WT1, DMRT1 and SOX9 were overexpressed (Fig. 3a, b). Notably, only the addition of GATA4, but not the addition of the three other TFs, to the reprogramming combination increased the percentage of AMH:EGFP+ cells. Several TFs, including NR5A1 and GATA4, are key regulators establishing the mouse Sertoli cell identity ^6^. Therefore, unsurprisingly, NR5A1 and GATA4 were sufficient to reprogram human fibroblasts to Sertoli-like cells. WT1 and SOX9 are also determining factors in mouse Sertoli cells, but their reprogramming abilities were lower than those of NR5A1 and GATA5. SOX9 interacts with NR5A1 to trigger the specific onset of AMH expression ^47^, but SOX9 overexpression in our reprogramming experiments only slightly increased the percentage of AMH:EGFP+ cells in a few combinations of TFs (Fig. 3a, b).

The cellular characterizations of the 2F-hiSCs showed that these cells carried many known properties of Sertoli cells. Lipid metabolism in Sertoli cells is important for providing nutritional and energy supplies to germ cells ^42^. Previous studies have shown that the high metabolic activity requirements for lipid and steroid synthesis are associated with the differentiated status of Sertoli cells ^48,49^. Notably, the hiSCs in the present study exhibited a high expression of lipid and steroid related genes (Fig. 3d), and the WT hiSCs exhibited a higher expression than the CX43KO hiSCs (Fig. 7h). These results suggest that the hiSCs were similar to differentiated Sertoli cells *in vivo*. The lipid droplet staining assays clearly showed the presence of high numbers of lipid droplets in the hiSCs but not the fibroblasts used for reprogramming (Fig. 4b).

Sustaining the viability and differentiation of spermatogonia cells is the most essential function of Sertoli cells. We examined the ability of hiSCs to sustain mouse spermatogonia cells because of the inaccessible procurement of human spermatogonia cells from hospitals. Previous studies performing xenotransplantation experiments transplanting primate spermatogonia stem cells into mouse testis demonstrated that primate germ cells colonized in the seminiferous tubules of the recipient mice, but the primate germ cells did not differentiate because of the evolutionary differences between these species^50^. The isolated mouse spermatogonia cells in our study attached and survived on the hiSCs, but no differentiated spermatocytes were detected (Fig. 5b, c; data not shown). These results suggest that the reprogrammed Sertoli cells were only capable of providing a niche for the survival of male germ cells *in vitro*. Recent studies have reported *in vitro* derivations of immature human gametes from hESCs ^16,17,51^. The male and female gametes in these studies were immature likely due to the absence of fully competent somatic cells in the differentiating culture. Therefore, adding induced Sertoli cells or granulosa cells to *in vitro* differentiating cultures may be a key step in obtaining fully functional gametes *in vitro*.

Another prominent role of Sertoli cells is the creation of an immune-privileged environment to protect germ cells from immune attacks of lymphocytes. Many studies have documented that Sertoli cells secrete factors that suppress the proliferation of T cells, B cells and NK cells ^14,45^, including the suppression of IL-2 from T cells^52^. Our results indicate that culture media incubated with hiSCs suppress the proliferation of Jurkat cells (the immortalized cell line of human T lymphocytes) and reduce IL-2 production in Jurkat cells treated with hiSC-conditioned media (Fig. 6b,c). Many genes previously known to participate in immune effector processes were upregulated in the hiSCs compared to those in fibroblasts (Fig. 6d), suggesting that hiSCs modulate immune suppression via multiple immunological pathways. Notably, all 293FT cells co-transplanted with hiSCs into two different immunocompetent mice survived longer than their counterpart control cells co-transplanted with fibroblasts (Fig. 6e and Supplementary Fig. 5). These results suggest that the hiSCs protected the xenotransplanted 293FT cells from host immune cells. Previous studies have reported that Sertoli cells protect many cell types in allogenic and xenogenic transplantation, and several studies used immunosuppressive drugs or immune-deficient mice in long-term follow-up experiments ^11,12^. One study xenotransplanted porcine Sertoli cells with rat islet cells into the kidney capsule of immunocompetent rats^53^. The islet allografts survived 8 to 9 days when 1.5 × 10^6^ porcine Sertoli cells were co-transplanted. In our experiments, 2.5 × 10^5^ hiSCs were co-transplanted with 293FT cells and survived at least 10 days in two mice. Therefore, our xenotransplanted 293FT cells survived a similar or longer duration with lower numbers of reprogrammed human Sertoli cells in immunocompetent mice. Long-term transplantations of hiSCs with clinically relevant cell types, including pancreatic islets and skin grafts, should be investigated for potential clinical applications, such as the treatment of diabetes and skin burns. Our hiSCs have the advantage of being of human origin, which alleviates the issue of xenotransplantation and potential of animal virus infection.

Reprogrammed Sertoli cells may be used as a model for examining the cellular and genetic mechanisms of human Sertoli cell biology. A previous study reported an association between Sertoli-only syndrome (SCO) and the lower mRNA expression of CX43 ^32^ and suggested that the absence of CX43 rendered the Sertoli cells more immature. However, the precise mechanism by which the absence of CX43 causes infertility is unclear. Our studies revealed that multiple pathways, including lipid metabolism and nucleobase catabolism, were more highly expressed in the WT than in the CX43KO hiSCs, indicating that the absence of CX43 disrupts these molecular pathways in Sertoli cells. The expression of several markers of immature Sertoli cells was higher and the expression of markers of mature Sertoli cells was lower in CX43KO than in the WT hiSCs. Taken together, our results suggest that the deletion of CX43 disrupts multiple molecular pathways and delays the maturation of Sertoli cells.

## AUTHOR CONTRIBUTIONS

J.L., N.W., J.D., Y.G. and L.L. carried out all the experiments; J.H. and J.L. conducted all bioinformatics analysis, J.L. and K.K. designed experiments and wrote the manuscript.

## ACKNOWLEDGMENTS

We are indebted to Prof. Zuping He for giving us the primary human Sertoli cells. Research funding is provided by the Ministry of Science and Technology of China [2018YFA0107703; 2017YFC1001601]; and Cross-discipline Foundation of Tsinghua University.

## Methods

### Human ES cell culturing and fibroblast differentiation

Human ES cell line used in this study were H1 (XY) and H9 (XX), purchased from WiCell, Inc. Undifferentiated H1 and H9 were maintained on MEF feeder cells as previous described^1^. All cells were cultured at 37°C in a humidified incubator supplied with 5% CO_2_. ES medium were standard knockout serum replacer (KSR) consisted of 20% knockout serum replacer, 0.1mM nonessential amino acids, 1mM L-glutamine, 0.1mM -mercaptoethanol, and 4 ng/ml recombinant human basic fibroblast growth factor (bFGF, R&D systems).

To obtain adherent fibroblast differentiation, human ESC clones were first transferred to 1% Matrigel coated plates by colony picking using a glass needle and cultured in ES cell conditioned medium (ES medium incubated overnight on irradiated MEFs) for ~5 days to remove all the residual MEF cells. Differentiation of hESCs to fibroblasts (dH1) began after aspirating of conditioned media, washing with PBS without Ca^2+^ and Mg^2+^ twice, and replacing with differentiation media (knockout DMEM with 20% fetal bovine serum, 0.1mM nonessential amino acids, 1mM L-glutamine). Differentiation media was changed every 3 days. When the cell reached 90% confluency (~7 days), the resulting dH1 were collected and kept in liquid nitrogen tank for long time preservation or passaged by the ratio 1:3 if more cells was needed.

### Human pulmonary fibroblasts and adult Sertoli cells culturing

Human pulmonary fibroblasts (HPF) were purchased from National Infrastructure of Cell Line Resource. Human adult Sertoli cells were a generous gift from Dr. Zuping He of Shanghai Jiao Tong University. The HPF cells were maintained at 75 ml culture flask in 15 ml DMEM culture medium (Corning) containing 10% FBS (Gibco), 0.1 mM nonessential amino acids and 1 mM L-glutamine. As for primary adult Sertoli cells, they were procured and cultured as previously described ^2^ in DF12 medium consisting of DMEM/F12 and 10% FBS (Gibco) and passaged every 5 days when cell confluence reached 80%. All of the cells above were cultured in a 37 °C humidified incubator supplied with 5% CO_2_.

### Overexpression vector construction and lentivirus Production

All overexpression vectors carrying EF1α promoter and desired gene were constructed using the Gateway system (Invitrogen) as previously described^1^. Briefly, the candidate cDNA was first introduced into pENTR/1A or pENTR/D-topo donor vectors and transferred to 2K7 destination vectors with EF1α promoter in pENTR/5’ topo by LR recombination ^3^. Modified destination plasmids containing the cDNA were then introduced into 293FT cells together with the helper plasmids vsvg and Δ8.9 by LIPO3000 (Invitrogen) transfection to produced virus. Approximately, a total of 37 ml of virus supernatant was harvested on day1 and day3 after transfection and filtered with a 0.45μm filter. At the time of virus infection, 8 μg/ml of polybrene was supplemented to increase infection efficiency.

### AMH:EGFP reporter and creating reporter cell line

1.6 kb of human AMH promoter was PCR amplified from genomic DNA of 293FT cells and cloned to pENTR5’-TOPO. Cloned plasmids were then recombined with pENTR/D-TOPO that carried the EGFP cDNA to create p2K7-AMH:EGFP recombinant plasmid and generated lentiviral supernatant as described above in overexpression vector construction section. Fibroblast HPF or dH1 in early passage (with 50% confluence) were transduced overnight on plate in fibroblast medium and recovered for one day after removal of virus. Subsequent drug selection by blasticidin (10 μg/ml) required another 3 days. Selected human fibroblasts were passaged for two times to expand the cell number for further experiments or frozen in liquid nitrogen.

### hiSC reprogramming and enrichment by FACS

Sertoli-like cell reprogramming was carried out in flask. In brief, human fibroblast cells carrying AMH reporter were seeded 1 day prior to overnight transduction with lentivirus NR5A1, GATA4, DMRT1, SOX9 and WT1 (for 5F-hiSCs) or NR5A1 and GATA4 (for 2F-hiSCs), recovered for 24 hours, following by drug selection with geneticin (1 mg/ml) for 5 days. After drug selection, transduced cells were cultured in hiSCs medium, maintained for the indicated reprogramming duration, and harvested for FACS by digestion with TrypLE Express (Invitrogen). The single cell suspension for FACS was prepared with MACS medium (10% FBS in PBS with 0.0125 mM EDTA) and filtrated through a BD cell sorter. Cell sorting was proceeded on a high-speed cell sorter (Influx, BD) and was sorted to collecting tube containing DF12 medium. Experiment aiming at examining different combinations of reprogramming factors was performed in 6 well plates and all possible combinations of NR5A1, GATA4, DMRT1, SOX9 and WT1 were tested (See table S2).

### Quantitative PCR and statistical analysis

Quantitative PCR was conducted as previously described ^1^. Briefly, total RNA was collected according to the instructions provided by QIAGEN RNeasy kit (QIAGEN) or TRIZOL (Invitrogen). CDNA was generated by EasyScript One-Step gDNA Removal and cDNA Synthesis SuperMix kit (TRANSGEN) according to manufacturer’s protocols using up to 500 ng RNA for each sample. 20 μl reaction (for Bio-rad 96-well System) were prepared and conducted with TransStart Green qPCR SuperMix kit (TRANSGEN). Gene expression was calculated using Bio-Rad CFX Manager program for relative expression formulation (dC(t)) and normalized to housekeeping genes (ACTB or GAPDH). Then, the gene expression of different samples were again normalized to the expression of control cells infected by p2k7 empty virus (CTRL) and reported as fold change ^4^. Statistical analysis was carried out using Student’s t-test or one-way ANOVA by Prism 6.0 software.

### RNA sequencing

~2.5 × 10^5^ cells were collected by FACS and total RNA was extracted by TRIZOL (Invitrogen). The quality and integrity of the purified RNAs was checked by Agilent 2100 bioanalyzer. Qualified RNA from the following samples was used for RNA Sequencing analysis: (1) HPF or dH1 carrying AMH:EGFP reporter, transduced with empty virus p2k7 and followed the same reprogramming procedure as described above. (2) Day10 hiSCs generated with 5 factors (5F-hiSCs) or generated from 2 factors (2F-hiSCs). (3) Human primary adult Sertoli cells (aSCs) cultured *in vitro* in DMEMF12 medium + 10% FBS. Sequencing libraries preparation and sequencing operations were carried out by ANNOROAD, a company providing RNA sequencing service, complied with the whole set of processes from Illumina.

### RNA-seq data processing

All RNA-seq data were mapped to human genome build hg19 (UCSC) by TopHat (version 2.1.1), reads from PCR duplicates were dropped. The gene expression level was calculated by Cufflinks (version 2.2.1) using the protein coding genes GTF file extracted from Ensembl database (Homo_sapiens.GRCh37.75.gtf). Read counts were obtained using HTSeq (version 0.9.1). Differentially expressed genes (DEGs) were analyzed using R package DESeq2 and selected using p-value < 0.01 as a threshold. Heat map was plotted using heatmap.2 function of R, and gene expressions were scaled to FPKM or Z-scores. We used R function cor to do sample correlation clustering and K-means clustering was performed using Cluster 3.0 package (K = 3, Spearman Correlation, Complete-linkage) and clustered heat maps were produced by TreeView.

### Immunofluorescence of cultured cells

Dissociated cells from FACS enrichment were collected onto a slide by Cytospin (800 r.p.m. for 5 min) or replated onto a 6-well plate. After that, cells were washed 1 time with PBS, fixed in 4% paraformaldehyde for 10 minutes and treated with 2.5% Triton X-100 for 15 minutes. For antibodies staining, slides were first blocked in 2.5% donkey serum for 1 hour, then incubated overnight in 4°C with primary antibody (1:200 for KRT18, 1:100 for NuMA, 1:200 for CX43, 1:100 for VASA, all rabbit-derived, Abcam; 1:50 for DAZL, mouse-derived, Bio-Rad; 1:100 for NR5A1, GATA4, WT1, SOX9, DMRT1, all rabbit-derived, Proteintech). Slides were then washed five times (each 3 minutes) with PBST (0.1% Tween-20/PBS), followed by anti-rabbit secondary antibodies (anti-rabbit-555 or anti-rabbit-488, Invitrogen) incubating for 1 hour at room temperature and washing for another 5 times. All sections were then mounted with Prolong Diamond Anti-Fade Mounting Reagent (ThermoFisher) and cover slip.

### Effect of 2F-hiSCs conditioned medium on Jurkat cell proliferation and IL-2 production

1 ml conditioned medium were collected from 3.5 × 10^4^ of 2F-hiSCs or dH1-2K7 fibroblast (control) cultured at 50% confluence 48 h after plating. For each assay, 2 × 10^4^ human T lymphocytes (Jurkat E6-1 cells, gift from Professor Hai Qi, Tsinghua University) were seeded in 1 well of 96-well plate with 120 μl 1064 medium with 10% FBS, 0.1mM nonessential amino acids, 1mM L-glutamine, 0.1mM–mercaptoethanol and added with indicated amount of conditioned medium from either 2F-hiSCs or dH1-2K7 (as control). The Jurkat cells were then cultured in a 37°C incubator with 5% CO_2_ for 3 days. Metabolism of WST-1 was used to determine the proliferative ability of lymphocytes in each well according to manufacturer’s instructions (Beyotime, Co., Ltd.). Three hours prior to analysis, 12 μl of WST-1 was added to each well and the final absorbance was measured by a microplate reader at 450 nm (SPECTRA max PLUS, Molecular Devices Inc.). 1064 medium alone with WST-1 was used as a control to subtract background absorbance. For IL-2 quantification, 2 × 10^5^ of Jurkat cells were seeded in one 6-well plate containing either 50% 2F-hiSCs or dH1-2K7 conditioned medium and cultured for 3 days. Then, cells were collected and lysed with 20 μl RIPA buffer. The concentration of IL-2 was determined by ELISA kit according to the manufacturer’s instructions (Cusabio Biotech, Co., Ltd.).

### Cell migration assays

Confluent P8 HUVEC cells (a gift from Professor Jie Na, Tsinghua university) were cultured with fresh ECM medium (Sciencell) overnight before experiment. Then cells were incubated with medium containing 2.5 μM Calcein-AM fluorescent dye for one hour in incubator supplied with 37°C, 5% CO_2_. Cells were then trypsinized, counted and suspended in migration medium (50% ECM+50% MEF medium). Migration assays were carried out in Corning FluoroBlok 24-multiwell insert plate with 8.0 μm pores (Cat. No. 351157. Corning). Prior to seeding HUVEC (150 k, in 100 μl volume) into the insert, 50 % conditioned medium pre-incubated with hiSCs or dH1 was mixed with 50% ECM and added to the basal chamber (600 μl in total). Following incubation of HUVECs at 37°C, 5% CO2 needed for 20 hours. The migration cells passed through the membrane were monitored by Calcein-AM with the help of an inverted microscope (Leica™) at 485/535 nm (Ex/Em). Images were captured using Leica-Pro software provided by the microscope company. The number of migration cell was measured by calculating of the Calcein-AM green signal of each image.

### Mouse PGC isolation and coculture with 2F-hiSCs

Mouse spermatogonia cells were isolated from ~24 testis of C57BL/6 mice at day 6 after birth according to previous protocol with minor modifications ^5^. Briefly, the decapsulated testis and seminiferous tubules were detached with the help of tweezers. After washing with DPBS for 3 times, the seminiferous tubules were transferred to a new 15 ml tube and subjected to enzymatic digestion. First, the seminiferous tubules were digested with 1mg/ml collagenase IV for 5 min at 37°C in a water bath with shaking, then centrifuged at 50× *g* to collect the segregated tubules and washed 3 times with DMEM/F12 medium. Completed disintegration of the tubules were achieved by treating with 0.25% trypsin for 5 min at 37°C with gentle shaking and pipetting. The suspension, containing primarily Sertoli cells, peritubular cell and male germ cells, was then pelleted at 500× *g* for 5 min at room temperature and washed once. Finally, cells were resuspended in 2 ml culture medium composed of DMEM/F12 medium with 10% FBS and seeded on 2 well of 6-well plates (Corning) and incubated at incubator supplied with 37°C, 5% CO_2_. After attachment for 24 hours, the somatic cells were tightly attached to the dish and formed patches, while some of the spermatogonia cells were just loosely attached on the somatic cells. 24 hours prior to coculture experiments, ~1.5 × 10^5^ dH1 or 2F-hiSCs were plated to 1 well of the 48 well plate. The spermatogonia cells were collected by gently washing with fresh media and transferred to the well plated with the dH1 or 2F-hiSCs in DMEMF/F12 medium with 10% FBS.

### Co-transplantation of hiSCs with 293FT cells

All animal experiments were approved by the Institutional Animal Care and Use Committees at Tsinghua University. ~1.3 × 10^6^ number of 293FT cells stably transduced with a luciferase reporter were mixed with 2.5 × 10^5^ cells of either dH1 or 2F-hiSCs in 100 μl Matrigel. The suspension was transplanted into C57B6/6J (Purchased from Charles River Laboratories) mice with normal immune system by subcutaneous injection. Live imaging of transplanted mice was performed at the indicative day after transplantation. 15 minutes prior to live imaging, 100 μl of D-luciferin (15mg/ml, 10 μL/g mouse weight) was injected to the mice by intraperitoneal injection and luciferase activity was measured in the IVIS Spectrum machine (PerkinElmer Health Sciences, Inc.).

